# TCR meta-clonotypes for biomarker discovery with tcrdist3: identification of public, HLA-restricted SARS-CoV-2 associated TCR features

**DOI:** 10.1101/2020.12.24.424260

**Authors:** Koshlan Mayer-Blackwell, Stefan Schattgen, Liel Cohen-Lavi, Jeremy Chase Crawford, Aisha Souquette, Jessica A. Gaevert, Tomer Hertz, Paul G. Thomas, Philip Bradley, Andrew Fiore-Gartland

## Abstract

As the mechanistic basis of adaptive cellular antigen recognition, T cell receptors (TCRs) encode clinically valuable information that reflects prior antigen exposure and potential future response. However, despite advances in deep repertoire sequencing, enormous TCR diversity complicates the use of TCR clonotypes as clinical biomarkers. We propose a new framework that leverages antigen-enriched repertoires to form meta-clonotypes – groups of biochemically similar TCRs – that can be used to robustly identify and quantify functionally similar TCRs in bulk repertoires. We apply the framework to TCR data from COVID-19 patients, generating 1831 public TCR meta-clonotypes from the 17 SARS-CoV-2 antigen-enriched repertoires with the strongest evidence of HLA-restriction. Applied to independent cohorts, meta-clonotypes targeting these specific epitopes were more frequently detected in bulk repertoires compared to exact amino acid matches, and 59.7% (1093/1831) were more abundant among COVID-19 patients that expressed the putative restricting HLA allele (FDR < 0.01), demonstrating the potential utility of meta-clonotypes as antigen-specific features for biomarker development. To enable further applications, we developed an open-source software package, *tcrdist3*, that implements this framework and facilitates flexible workflows for distance-based TCR repertoire analysis.

## INTRODUCTION

An individual’s unique repertoire of T cell receptors (TCRs) is shaped by antigen exposure and is a critical component of immunological memory, contributing to recall responses against future infectious challenges (Emerson et al., 2017; Welsh and Selin, 2002). With the advancement of immune repertoire profiling, TCR repertoires are a largely untapped source of biomarkers that could potentially be used to predict immune responses to a wide range of exposures including viral infections (Wolf et al., 2018), tumor neoantigens (Ahmadzadeh et al., 2019; Chiou et al., 2021; Kato et al., 2018), or environmental allergens (Cao et al., 2020). The TCR repertoire is characterized by its extreme diversity, originating from the genomic V(D)J gene recombination of receptors in development. Between 10^9^-10^10^ unique clonotypes - T cells with distinct nucleotide-encoded receptors - are maintained in an adult human TCR repertoire (Lythe et al., 2016). The diversity, both within and between individuals, presents major hurdles to biomarker development. Researchers have used antigen-enrichment of T cell repertoires (e.g. peptide-major histocompatibility complex (MHC) tetramer sorting) to focus on TCR diversity of relevant targets, however this experimental strategy, which depends on knowing the peptide antigen and it’s MHC restriction reveals the breadth of potential TCRs able to recognize even a single antigen (Coles et al., 2020; Meysman et al., 2019), which complicates detection of population-wide signatures of antigen exposure. Indeed, mathematical modeling suggests that only 10-15% of single chain TCRs are public or shared frequently by multiple individuals (Elhanati et al., 2018), which is consistent with observations from extremely deeply sequenced human repertoires (Soto et al., 2019). Despite advances in high-throughput next-generation TCR amplicon sequencing, only a fraction of the repertoire can be assayed, making it difficult to reproducibly sample many relevant TCR clonotypes from an individual, let alone reliably detect public clonotypes in a population. In practice, the problem is exacerbated by unequal sampling depth. Thus, individual T cell clonotypes are currently sub-optimal and under-powered for population-level investigations of TCR specificity, which limits their application in the development of TCR-based clinical biomarkers.

In this study we used antigen enriched TCR repertoires to form “meta-clonotypes”: groups of TCRs with biochemically similar complementarity determining regions (CDRs) that likely share antigen recognition. Meta-clonotypes were implemented using a centroid TCR sequence and a biochemical radius that determines whether other TCRs are sufficiently similar to be grouped together; the appropriate radius was determined by comparing the proportion of similar TCRs in antigen-enriched and unenriched data. A CDR3 “motif” is also constructed from the TCRs within the radius, which further refines the specificity of meta-clonotype definition. Together the radius and the motif can be used to search for conformant TCRs in large bulk-sequenced repertoires and quantify their abundance (Figure 1). We find that TCR centroids, which are often private, gain publicity as meta-clonotypes.

**Figure 1.**
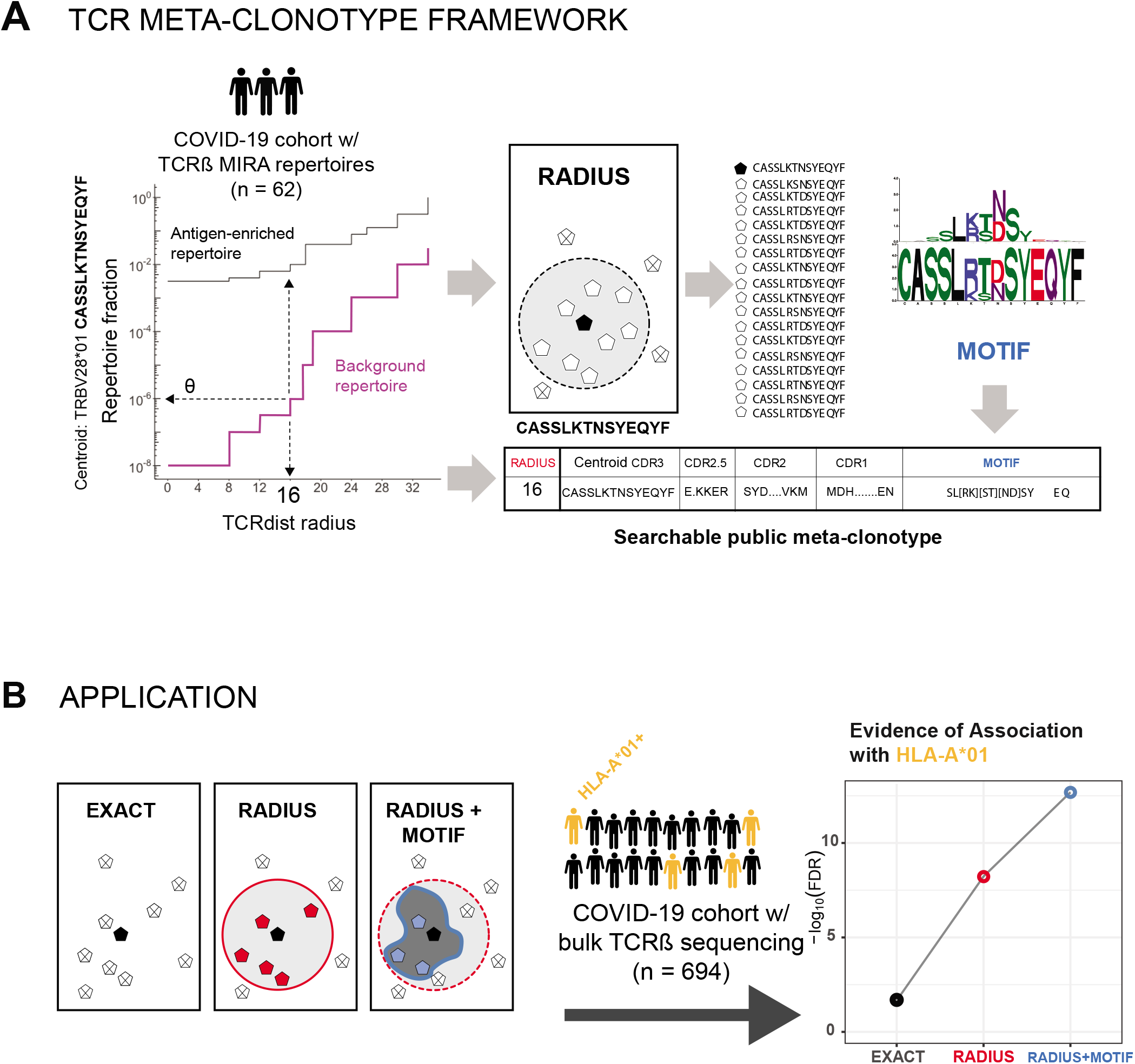
TCR meta-clonotype framework and application. (A) Framework: antigen-enriched repertoires were used together with antigen-unenriched background repertoires to engineer TCR meta-clonotypes that define biochemically similar TCRs based on a centroid TCR and a TCRdist radius. Antigen-enriched TCRs came from CD8+ T cells activated by SARS-CoV-2 peptides that were previously discovered (Nolan et al., 2020) in 62 individuals diagnosed with COVID-19 using MIRA (Multiplex Identification of Antigen-Specific T Cell Receptors Assay, Klinger et al., 2015). With each clonotype from the antigen-enriched TCRs, we used *tcrdist3* to evaluate the repertoire fraction spanned at different TCRdist radii within (i) its antigen-enriched repertoire (black) and (ii) a control V- and J-gene matched, inverse probability weighted background repertoire (purple). The set of antigen-enriched TCRs spanned by the optimal radius were then used to develop an additional meta-clonotype motif constraint based on conserved residues in the CDR3 (see Methods for details). An example logo plots shows the CDR3 β-chain motif formed from TCRs – activated by a SARS-CoV-2 peptide (MIRA55 ORF1ab amino acids 1316:1330, ALRKVPTDNYITTY) – within a TCRdist radius 16 of this meta-clonotype’s centroid. (B) Application: TCR meta-clonotypes were used to quantify the frequency of putative SARS-CoV-2 antigen-specific TCRs in a large diverse cohort, from whom bulk TCR repertoires were collected 0-30 days after COVID-19 diagnosis (n=694). Meta-clonotypes were evaluated based on their association with a restricting HLA allele. In most cases, evidence of HLA-restriction was stronger for meta-clonotypes (RADIUS or RADIUS+MOTIF) compared to using exact matches to the centroid TCR (EXACT), demonstrated by lower false-discovery rate (FDR) adjusted q-values and larger HLA regression coefficients in beta-binomial count regression models that account for sequencing depth and control for patient age, sex, and days from diagnosis.

The expanded publicity of meta-clonotypes provides an opportunity to develop population-level biomarkers of clinical outcomes that depend on antigen-specific features of the TCR repertoire, such as disease severity in natural infection or the level of vaccine-induced protection. Shifting the focus of repertoire analysis from clonotypes to meta-clonotypes increases statistical power by reducing the inherent sparsity of finite repertoire samples and increasing the precision with which antigen-specific cell abundance can be estimated. A number of existing tools enable grouping of TCRs by sequence similarity (Table S1); for example, VDJtools (TCRNET) and ALICE evaluate networks of similar TCR β- or TCR α-chain CDR3s based on a maximum edit-distance of one amino acid substitution, insertion or deletion, while GLIPH2 groups similar TCRs based on shared amino acid k-mers in identical length CDR3s (Glanville et al., 2017; Huang et al., 2020; Pogorelyy et al., 2019; Pogorelyy and Shugay, 2019; Ritvo et al., 2018; Shugay et al., 2015). Previously, we introduced TCRdist, a weighted multi-CDR, biochemically informed distance metric that enabled grouping of paired αβ TCRs by antigen specificity, based on their sequence similarity (Dash et al., 2017). Here, we describe a new application of TCRdist that guides formation of meta-clonotypes optimized for biomarker development. This application is made possible by a new open-source Python3 software package *tcrdist3* that brings new flexibility to distance-based repertoire analysis, allowing customization of the distance metric, analysis of γδ TCRs, and at-scale computation with sparse data representations and parallelized, byte-compiled code.

Here we first describe a novel analytical framework for identifying meta-clonotypes in antigen-enriched repertoires. The framework is then applied to a large publicly available dataset of putative SARS-CoV-2 antigen-associated TCRs with the objective of identifying meta-clonotypes that could be used as features in further developing SARS-CoV-2 related biomarkers (Figure 1). One of the distinguishing characteristics of SARS-CoV-2 infection is the wide range of potential exposure outcomes, from transient, asymptomatic infection to severe disease requiring hospitalization and intensive care. While there are high quality biomarkers for detecting active SARS-CoV-2 infection via viral RNA qPCR (Nalla et al., 2020) and prior exposure via antibody ELISA (Espejo et al., 2020), additional biomarkers capable of predicting susceptibility to symptomatic infection or severe disease could help guide clinical care and public health policy. Several studies have begun to describe the cellular adaptive immune responses that are elicited by SARS-CoV-2 infection and how they correlate with disease severity (Grifoni et al., 2020; Le Bert et al., 2020; McMahan et al., 2020; Tan et al., 2021; Wang et al., 2020; Weiskopf et al., 2020). These and other studies have also established that 20-50% of unexposed individuals have T cell responses to SARS-CoV-2, raising the hypothesis that prior exposure to “common-cold” coronaviruses or other viral antigens may shape the response to SARS-CoV-2 infection (Sette and Crotty, 2020; Welsh and Selin, 2002). T cells likely play an integral role in SARS-CoV-2 pathogenesis and may be an important target for biomarker development. For instance, a TCR biomarker of pre-existing SARS-CoV-2 responses could help predict the course of infection. A T cell-based biomarker might also play a role in vaccine development, for which immunological surrogates of vaccine-induced protection or response durability are highly valued. Most published studies have had limited ability to determine quantitative immunodominance hierarchies, relying on pooled peptide assays, due to the large size of the SARS-CoV-2 proteome and HLA diversity; direct repertoire measurement tied to identified epitopes is a complementary approach to resolve the associated magnitude and specificity of the total T cell response.

One recent study to elucidate the role of cellular immune responses in acute SARS-CoV-2 infection examined the T cell receptor repertoires of patients diagnosed with COVID-19 disease. Researchers used an assay based on antigen stimulation and flow cytometric sorting of activated CD8+ T cells to sequence SARS-CoV-2 peptide-associated TCR β-chains; the assay is called “multiplex identification of T-cell receptor antigen specificity” or MIRA (Klinger et al., 2015). Data from these experiments were released publicly in July 2020 by Adaptive Biotechnologies and Microsoft as part of “immuneRACE” and their efforts to stimulate science on COVID-19 (Nolan et al., 2020; Snyder et al., 2020). The MIRA antigen enrichment assays identified 269 sets of TCR β-chains associated with CD8+ T cells activated by exposure to SARS-CoV-2 peptides, with TCR sets ranging in size from 1 - 16,607 TCRs (Table S1). The deposited immuneRACE datasets also included bulk TCR β-chain repertoires from 694 patients within 0-30 days of COVID-19 diagnosis. To demonstrate potential uses of our new analytical tools for TCR repertoire analysis and to accelerate understanding of the cellular responses to SARS-CoV-2 infection, we present analyses of these data with a focus on an integration of the peptide-associated MIRA TCR repertoires with bulk repertoires from four COVID-19 observational studies that enrolled patients with diversity in age and geography (Alabama, USA n = 374; Madrid, Spain, n=117; Pavia, Italy, n=125; Washington, USA, n=78).

## FRAMEWORK

### Experimental antigen-enrichment allows discovery of TCRs with biochemically similar neighbors

Searching for identical TCRs within a repertoire - arising either from clonal expansion or convergent nucleotide encoding of amino acids in the CDR3 - is a common strategy for identifying functionally important receptors. However, in the absence of experimental enrichment procedures, observing T cells with the same amino acid TCR sequence in a bulk sample is rare. For example, in 10,000 β-chain TCRs from an umbilical cord blood sample, less than 1% of TCR amino acid sequences were observed more than once, inclusive of possible clonal expansions (Figure 2A). By contrast, a valuable feature of antigen-enriched repertoires is the presence of multiple T cells with identical or highly similar TCR amino acid sequences (Figure 2A). For instance, 45% of amino acid TCR sequences were observed more than once (excluding clonal expansions) in a set of influenza M1(GILGFVFTL)-A*02:01 peptide-MHC tetramer sorted sub-repertoires from 15 subjects (Dash et al., 2017). Enrichment was evident compared to cord blood for additional peptide-MHC tetramer sorted sub-repertoires obtained from VDJdb (Shugay et al., 2018), though the proportion of TCRs with an identical or similar TCR in each set was heterogeneous.

**Figure 2.**
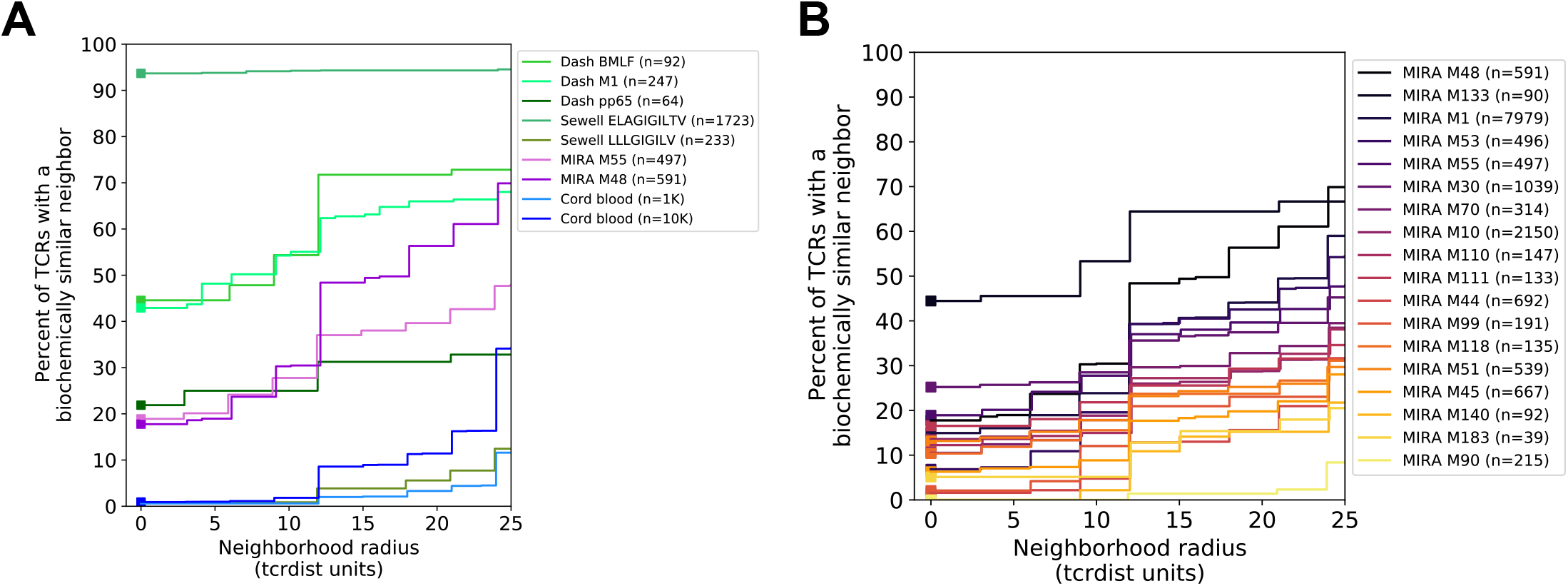
Experimental enrichment of antigen-specific TCRs. (A) TCR repertoire subsets obtained by single-cell sorting with peptide-MHC tetramers (data from Dash et al. and Sewell et al. via VDJdb; greens), MIRA peptide stimulation enrichment (MIRA55, MIRA48; purples), or random sub-sampling of umbilical cord blood (1,000 or 10,000 TCRs; blues). Biochemical distances were computed among all pairs of TCRs in each subset using the TCRdist metric. Neighborhoods were formed around each TCR using a variable radius (x-axis) and the percent of TCRs in the set with at least one other TCR within its neighborhood was computed. A radius of zero indicates the proportion of TCRs that have at least one TCR with an identical amino acid sequence (solid square). (B) Analysis of MIRA-enriched repertoires for which the participants contributing the TCRs were significantly enriched with a specific class I HLA allele (Table S5).

We investigated the degree to which the MIRA enrichment strategy employed by Nolan et al. (2020) identified TCRs with identical or similar amino acid sequences. In general, across multiple MIRA TCR β-chain antigen-enriched repertoires, the proportion of amino acid TCR sequences observed more than once was generally lower than in the tetramer-enriched repertoires and varied considerably across the sets; some MIRA sets resembled tetramer-sorted sub-repertoires (Figure 2B; see MIRA133), while others were more similar to unenriched repertoires (Figure 2B; see MIRA90). The increased diversity in MIRA-enriched TCR sets versus tetramer-enriched TCR sets may, in part, be explained by: (i) peptides being presented by the full complement of the native host’s MHC molecules compared to a single defined peptide-MHC complex, (ii) recruitment of lower affinity receptors, or (iii) non-specific “bystander” activation in the MIRA stimulation assay.

### TCR biochemical neighborhood density is heterogeneous in antigen-enriched repertoires

We next investigated the proportion of unique TCRs with at least one biochemically similar neighbor among TCRs with the same putative antigen specificity. We and others have shown that a single peptide-MHC epitope is often recognized by many distinct TCRs with closely related amino acid sequences (Dash et al., 2017); in fact, detection of such clusters in bulk-sequenced repertoires is the basis of several existing tools: GLIPH (Glanville et al., 2017; Huang et al., 2020), ALICE (Pogorelyy et al., 2019) and TCRNET (Ritvo et al., 2018). Therefore, to better understand new large-scale antigen-enriched datasets, like the SARS-CoV-2 MIRA data, we evaluated the TCR biochemical neighborhoods, defined for each TCR as the set of similar TCRs whose sequence divergence is within a specified radius. The radius was measured using a position weighted, multi-CDR TCR distance metric. Briefly, differences in the amino-acid sequences of the CDRs are totaled based on number of gaps (−4) and their BLOSUM62 substitution penalties (ranging from 0 to −4) with 3-fold weighting on CDR3 substitutions (see Methods for details of *tcrdist3* re-implementation of TCRdist); a one amino-acid mismatch in the CDR3 results in a maximal distance of 12 TCRdist units (tdus). As the radius about a TCR centroid expands, the number of TCRs it encompasses naturally increases; the rate of increase is more rapid in the antigen-enriched repertoires compared to the unenriched repertoires (Figure 2).

To better understand the relationship between the TCR distance radius and the density of proximal TCRs, we constructed empirical cumulative distribution functions (ECDFs) for each unique TCR found within a repertoire (Figure 3). The ECDF for each unique TCR (one line in Figure 3) shows the proportion of all TCRs within the indicated radius; those with sparse neighborhoods appear as lines that remain flat and do not increase along the *y*-axis even as the search radius expands. Moreover, the proportion of TCRs with sparse or empty neighborhoods (ECDF proportion < 0.001) is indicated by the height of the gray area plotted below the ECDF (Figure 3). We observed the highest density neighborhoods within repertoires that were sorted based on peptide-MHC tetramer binding. For instance, with the influenza M1(GILGFVFTL)-A*02:01 tetramer-enriched repertoire from 15 subjects, we observed that many TCRs were concentrated in dense neighborhoods, which included as much as 30% of the other influenza M1-recognizing TCRs within a radius of 12 tdus (Figure 3A). Notably there were also many TCRs with empty or sparse neighborhoods using a radius of 12 tdus (111/247, 44%) or 24 tdus (83/247, 34%). Based on previous work (Dash et al., 2017), we assume that the majority of these tetramer-sorted CD8+ T cells without many close proximity neighbors do indeed bind the influenza M1:A*02:01 tetramer. This suggests that TCRs within sparse neighborhoods represent less common modes of antigen recognition and highlights the broad heterogeneity of neighborhood densities even among TCRs recognizing a single pMHC.

**Figure 3.**
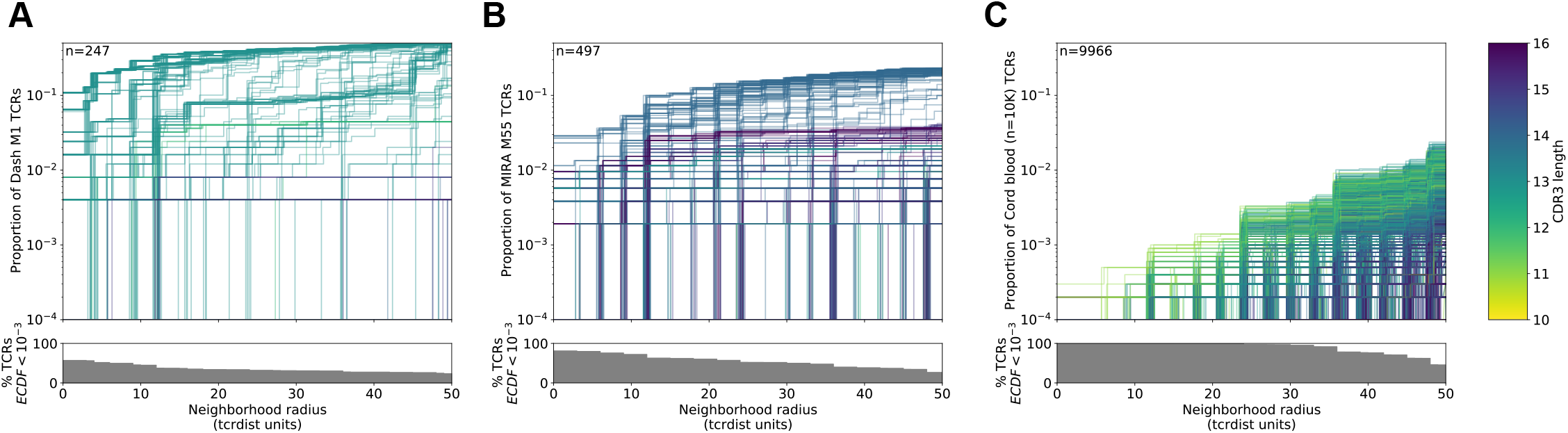
Heterogeneous TCR neighborhoods within experimentally antigen-enriched and unenriched repertoire subsets. TCR β-chains from (A) a peptide-MHC tetramer-enriched sub-repertoire, (B) a MIRA peptide stimulation-enriched sub-repertoire, or (C) an umbilical cord blood unenriched repertoire. Within each sub-repertoire, an empirical cumulative distribution function (ECDF) was estimated for each TCR (one line) acting as the centroid of a neighborhood over a range of distance radii (x-axis). Each ECDF shows the proportion of TCRs within the MIRA set with a distance to the centroid less than the indicated radius. ECDF color corresponds to the length of the CDR3-β loop. ECDF curves were randomly shifted by <1 unit along the x-axis to reduce overplotting. Vertical ECDF lines starting at 10^−4^ indicate no similar TCRs at or below that radius. Percentage of TCRs with an ECDF proportion < 10^−3^ (bottom panels), indicates the percentage of TCRs without, or with very few biochemically similar neighbors at the given radius.

Neighbor densities for individual TCRs within MIRA identified antigen-enriched repertoires were highly heterogeneous. Densities for an illustrative MIRA set are shown in Figure 4 (MIRA55:ORF1ab; 1316:1330 (amino acid); peptide ALRKVPTDNYITTY). Within this antigen-enriched repertoire, at 24 tdus 8.9% (44/497) of TCR neighborhoods included >10% of the other antigen-activated CD8+ TCRs (Figure 4A). As expected, TCR neighborhoods in the umbilical cord blood repertoire were sparser (Figure 4B); the densest neighborhood included only 0.13% of the repertoire at 24 tdus. We also noted that TCRs with empty neighborhoods tended to have longer CDR3 loops (Figure 4C). This is consistent with mathematical modeling approaches that show that TCRs with longer CDR3 loops have a lower generation probability (*P_gen_*) during genomic recombination of the TCR locus (Marcou et al., 2018; Murugan et al., 2012; Sethna et al., 2019). Absent strong selection for antigen recognition, TCRs with a low generation probability are thus likely to have a less dense biochemical neighborhood. Together, these observations suggest that biochemical neighborhood density is highly heterogeneous among TCRs and that it may depend on mechanisms of antigen-recognition as well as receptor V(D)J recombination biases (Thomas and Crawford, 2019).

**Figure 4.**
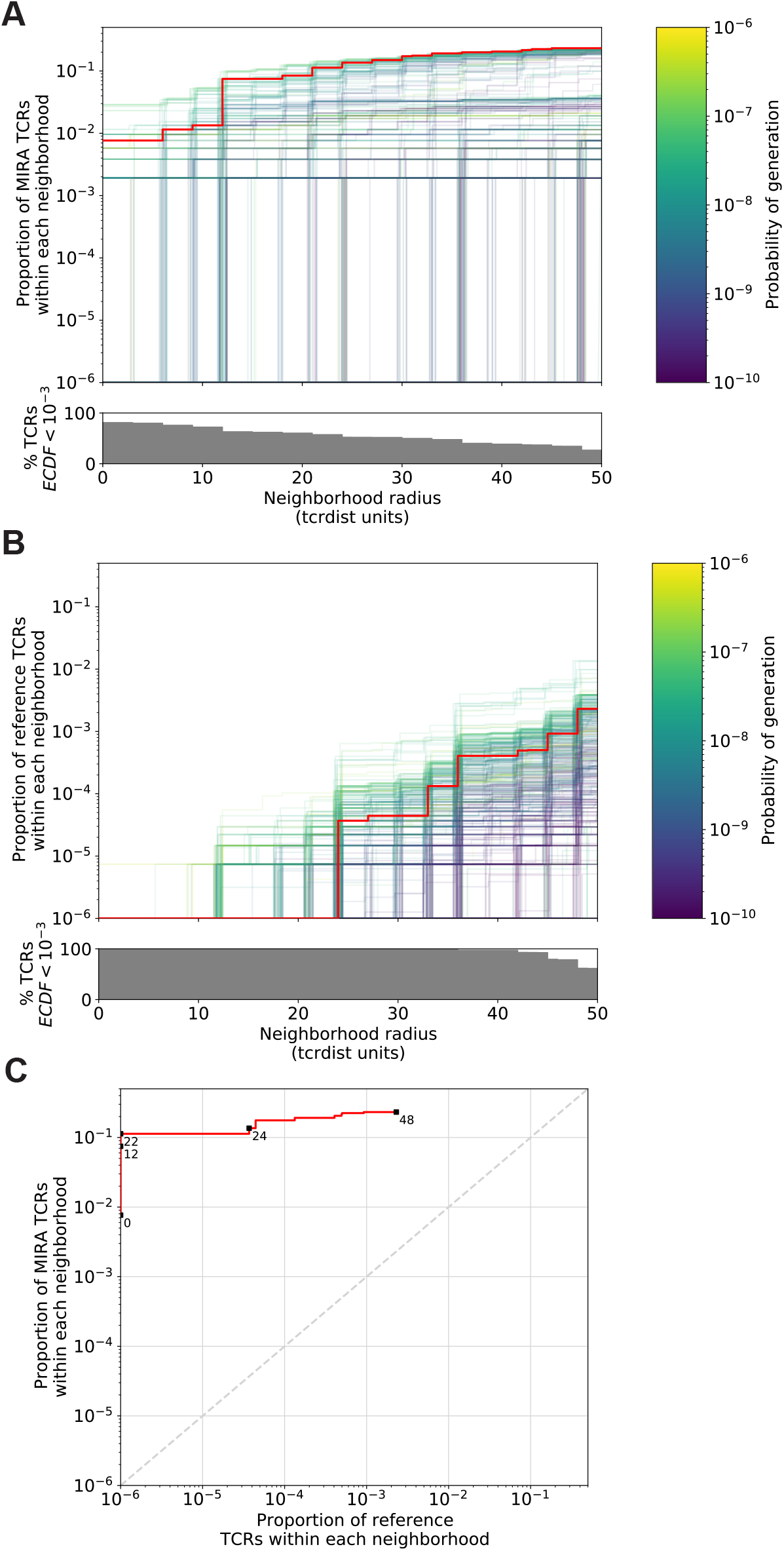
Radius-defined neighborhood densities within an antigen-enriched and a synthetic background repertoire. (A) Each TCR in the MIRA55 antigen-enriched sub-repertoire (one line) acts as the centroid of a neighborhood and an empirical cumulative distribution function (ECDF) is estimated over a range of distance radii (x-axis). Each ECDF shows the proportion of TCRs within the MIRA set having a distance to the centroid less than the indicated radius. The ECDF line color corresponds to the TCR generation probability (*P_gen_*) estimated using OLGA (Sethna et al., 2019). The ECDF curves are randomly shifted by <1 unit along the x-axis to reduce overplotting. The bottom panel shows the percentage of TCRs with an ECDF proportion < 10^−3^. (B) Estimated ECDF for each MIRA55 TCR based on the proportion of TCRs in a synthetic background repertoire that are within the indicated radius (x-axis). The synthetic background was generated using 100,000 OLGA-generated TCRs and 100,000 TCRs sub-sampled from umbilical cord blood; sampling was matched to the VJ-gene frequency in the MIRA55 sub-repertoire, with inverse probability weighting to account for the sampling bias (see Methods for details). (C) Antigen-enriched ECDF (y-axis) of one example TCR’s neighborhood (red line) plotted against ECDF within the synthetic background (x-axis). Example TCR neighborhood is the same indicated by the red line in (A) and (B). The dashed line indicates neighborhoods that are equally dense with TCRs from the antigen-enriched and unenriched background sub-repertoires.

### Meta-clonotype radius can be tuned to balance a biomarker’s sensitivity and specificity

The utility of a TCR-based biomarker depends on the antigen-specificity of the TCRs. Therefore, a key constraint on distance-based clustering is the presence of similar TCR sequences that may lack the ability to recognize the target antigen. To be useful, a meta-clonotype definition should be broad enough to capture multiple biochemically similar TCRs with shared antigen-recognition, but not excessively broad as to include a high proportion of non-specific TCRs, which might be found in unenriched background repertoires that are largely antigen-naïve. Because the density of neighborhoods around TCRs are heterogeneous, we hypothesize that the optimal radius defining a meta-clonotype may differ for each TCR. To find the ideal radius we proposed comparing the relative density of a radius-defined neighborhood in an antigen-enriched sub-repertoire (Figure 4A) to the density of the radius-defined neighborhood in an unenriched background repertoire (Figure 4B, 4C). This is similar to previous approaches taken by tools like ALICE and TCRNET, except that we employ a biochemically informed distance measure (TCRdist) and adjust the radius around each TCR to balance the antigen-enriched and unenriched neighborhood densities. The radius around each TCR defines a meta-clonotype that can be used to search for and quantify the abundance of conformant sequences in bulk repertoires (Figure 1A, 1B). For each TCR, its radius-defined meta-clonotype is more abundant within a repertoire and more prevalent in a population than the exact clonotype; for example, TCR meta-clonotypes formed from the MIRA55:ORF1ab TCR set were detected in 3 to 12 (median 6) of 15 HLA-A*01 participants in the MIRA cohort, despite 34 of the 46 centroid clonotype TCRs being private (i.e., found in only 1 of 15 HLA-A*01 participants). (Figure S1).

An ideal radius-defined meta-clonotype would include a high density of TCRs in antigen-experienced individuals indicative of shared antigen specificity, yet a low density of TCRs among an antigen-naïve background. We demonstrate this approach for selecting an optimal radius for TCRs in the MIRA55:ORF1ab dataset, which includes TCRs from 15 COVID-19 diagnosed subjects (see Methods for details about MIRA and the immuneRACE dataset). First, an ECDF is constructed for each centroid TCR showing the relationship between the meta-clonotype radius and its “sensitivity”: its inclusion of similar antigen-recognizing TCRs, approximated by the proportion of TCRs in the antigen-enriched MIRA set that are within the centroid’s radius (Figure 4A). Next, an ECDF is constructed for each TCR showing the relationship between the meta-clonotype radius and its “specificity”: its exclusion of TCRs with divergent antigen-recognition; this is assessed by computing the false-positive rate (one minus specificity) which is approximated by the proportion of TCRs in an unenriched background repertoire within the centroid’s radius (Figure 4B). Generating an appropriate set of unenriched background TCRs is important; for each set of antigen associated TCRs discovered by MIRA, we created a two part background. One part consisted of 100,000 synthetic TCRs whose TRBV- and TRBJ-gene frequencies matched those in the antigen-enriched repertoire; TCRs were generated using the software OLGA (Marcou et al., 2018; Sethna et al., 2019). The other part consisted of 100,000 umbilical cord blood TCRs sampled from 8 subjects (Britanova et al., 2017). This composition balanced denser sampling of sequences near the candidate meta-clonotype centroids with broad sampling of TCRs from an antigen-naïve repertoire. Importantly, we adjusted for the biased sampling by using the TRBV- and TRBJ-gene frequencies observed in the cord blood data (see Methods for details about inverse probability weighting adjustment). Using this approach, we are able to estimate the abundance of TCRs similar to a centroid TCR in an unenriched background repertoire of effectively ~1,000,000 TCRs, using a comparatively modest background dataset of 200,000 TCRs. While this may underestimate the true specificity since some of the neighborhood TCRs in the unenriched background repertoire may in fact recognize the antigen of interest, this measure is useful for prioritizing neighborhoods and selecting a radius for each neighborhood that balances sensitivity and specificity.

We find that the neighborhoods around TCR centroids with higher probabilities of generation consistently span a larger proportion of unenriched background TCRs across a range of radii, suggesting that a smaller radius may be desirable for forming meta-clonotypes from high *P_gen_* TCRs. With a large radius, most TCR centroids had high sensitivity and low specificity, indicated by the meta-clonotypes including both a high proportion of TCRs from the antigen-enriched and unenriched repertoires. Some TCRs had low sensitivity and specificity even at a radius of 24 tdus, indicative of a low *P_gen_* or “snowflake” TCR: a seemingly unique TCR in both the antigen-enriched and unenriched repertoires. However, radius-defined neighborhoods around many TCRs in the MIRA55:ORF1ab repertoire included 1 - 10% of the antigen-enriched repertoire (5-50 clonotypes) with a radius that included fewer than 0.0001% of TCRs (equivalent to 1 out of 10^6^) in the unenriched background repertoire, demonstrating a level of sensitivity and specificity that would be favorable for development of a TCR biomarker (Figure 4C, one example meta-clonotype).

## RESULTS

### Engineering meta-clonotype features for SARS-CoV-2

The MIRA antigen enrichment assays identified 269 sets of TCR β-chains associated with recognition of a SARS-CoV-2 antigen using CD8+ T cell enriched PBMC samples from 62 COVID-19 diagnosed patients. Of these, 252 included at least 6 unique TCRs from ≥ 2 individuals, which we refer to as MIRA1 - MIRA252 based on the number of sequences observed, in descending order (Table S2). From the MIRA enriched repertoires, all TCR clonotypes (defined by identical TRBV gene and CDR3 at the amino acid level) were initially considered as candidate centroids; only 2.7% of the clonotypes were found in more than one MIRA participant. For each candidate TCR, a meta-clonotype was engineered by selecting the maximum radius that limited the estimated number of neighboring TCRs in a bulk unenriched repertoire to less than 1 in 10^6^, estimated using an inverse probability weighted antigen-naïve background repertoire (see Methods). We then ranked the meta-clonotypes by their sensitivity approximated as the proportion of a centroid’s MIRA-enriched repertoire spanned by the search radius (diagrammed in Figure 1). Lower-ranked meta-clonotypes were eliminated if all included sequences were completely encompassed by a higher-ranked meta-clonotype; while this reduced redundancy, some overlap remained among meta-clonotypes. We further required that meta-clonotypes be public, including sequences from at least two subjects in the MIRA cohort. We found that 97 of the 252 MIRA sets (Table S6) had sufficiently similar TCRs observed in multiple subjects allowing formation of public meta-clonotypes. From 91,122 TCR β-clonotypes across these 97 MIRA sets -- targeting antigens in ORF1ab (n=35), S (n=27), N (n=10), M (n=7), ORF3a (n=7), ORF7a (n=4), E (n=2), ORF8 (n=2), ORF6 (n=1), ORF7b (n=1), and ORF10 (n=1) -- we engineered 4548 public meta-clonotypes, which spanned 15% (13,949/91,122) of the original TCR sequences (Table S6). The proportion of MIRA-enriched TCRs spanned by the meta-clonotypes ranged widely from <1% in MIRA25 to 63% in MIRA7, reflecting broad heterogeneity in the diversity of TCRs inferred as activated by each peptide in the assay.

As an example, the MIRA repertoire MIRA55 ORF1ab (TCRs associated with stimulation peptides ALRKVPTDNYITTY or KVPTDNYITTY) included 449 TCR clonotypes from 15 individuals. From the 449 potential centroids, we defined 40 public meta-clonotypes. Among these features, the radii ranged from 10-36 tdus (median 22 tdus), and the publicity - the number of unique subjects spanned by the meta-clonotype - ranged from 3 to 12 individuals (median 6). Meta-clonotype summary statistics for other enriched repertoires are provided in the Supplemental Materials (Table S6). The result was a set of non-redundant, public meta-clonotypes (Table S7, S8) that could be used to search for and quantify putative SARS-CoV-2-specific TCRs in bulk repertoires. In addition to the radius-defined meta-clonotypes (RADIUS), we also developed a modified approach that additionally enforced a sequence motif-constraint (RADIUS + MOTIF). The constraint further limited sequence divergence in highly conserved positions of the CDR3, requiring that candidate bulk TCRs match specific amino acids found in the meta-clonotype CDR3s to be counted as part of the neighborhood (see Figure 1 and Methods).

### Evidence of HLA-restriction in SARS-CoV-2 antigen-enriched sub repertoires

Given the integral role of HLA class I molecules in antigen presentation and TCR repertoire selection (DeWitt, 2018), we further focused on the 17 of the 269 MIRA sets that showed strong evidence of HLA-A or HLA-B restriction based on two criteria: (i) computational prediction of HLA binding to the SARS-CoV-2 stimulation peptides, and (ii) enrichment of an HLA among participants contributing MIRA TCRs. With each set of the MIRA TCRs and the associated SARS-CoV-2 peptides we used HLA binding predictions (NetMHCpan4.0) to identify the class I HLA alleles that were predicted to bind with strong (IC50<50 nM) or weak (50 nm< IC50 <500 nM) affinity to any of the 8, 9, 10, or 11-mers derived from the stimulation peptides (Tables S2, S3). For instance, the peptides associated with MIRA55 TCRs (ORF1ab amino acid positions 1316:1330) are predicted to preferentially bind A*01 (IC50 21 nM), B*15 (IC50 120 nM), and B*35 (IC50 32 nM), and peptides associated with MIRA51 TCRs (nucleocapsid amino acid positions 359:370) are predicted to bind A*03 (IC50 19 nM), A*11 (IC50 8 nM), and A*68 (IC50 9 nM).

Of the COVID-19 patient samples screened using the MIRA assay, HLA genotypes were available for 47 of 62 patients and only a subset of patients contributed TCRs to each of the MIRA sets. As a second indicator of HLA restriction, we tested whether the subgroup of patients contributing TCRs to each MIRA set was enriched with individuals expressing specific HLA class I alleles (2-digit resolution) (Table S5). We found that for 17 of the MIRA sets, the patients contributing TCRs were significantly enriched for at least one HLA-A or HLA-B allele (Fisher’s exact test p<0.001). In one case, all 13 A*01-positive, and only 2 of 34 A*01-negative, patients contributed to the MIRA55 TCR set (p=1e-7); as noted above, A*01 was also strongly predicted by NetMHCpan4.0 to bind the MIRA55 ORF1ab peptides. Similar patterns of enrichment and predicted binding were seen with A*01 expressing individuals and recognition of MIRA1:ORF1ab (HTTDPSFLGRY, p=1.9e-7) and MIRA45:ORF3a (SYFTSDYYQ, p=1.9e-7). Similarly, for the other 16 MIRA sets examined, we found consistent evidence between peptide binding prediction (IC50 < 500 nM) and MIRA participant genotype enrichment (fisher’s exact test p < 0.001) to support HLA-restriction (Table S5).

### HLA-associated abundance of SARS-CoV-2 meta-clonotypes in bulk repertoires of COVID-19 patients

We focused confirmatory analyses on TCR meta-clonotypes derived from the 17 SARS-CoV-2 MIRA-identified TCR sets that showed strongest evidence of HLA restriction by HLA-A or HLA-B alleles. We hypothesized that in an independent cohort of COVID-19 patients, the abundance of TCRs matching each meta-clonotype would be greater in patients expressing the restricting HLA allele. To test this hypothesis, we compared three TCR-based feature sets: (i) radius-defined meta-clonotypes, (RADIUS), (ii) radius and motif-defined meta-clonotypes (RADIUS+MOTIF) and (iii) centroid clonotypes alone, using TRBV-CDR3 amino acid matching (EXACT). Using the features in each set we screened TCRs from the bulk TCR β-chain repertoires of 694 COVID-19 patients whose repertoires were publicly released as part of the immuneRACE datasets (see Methods for details); these patients were not part of the smaller cohort that contributed samples for TCR identification in MIRA experiments. Testing the HLA restriction hypothesis required having the HLA genotype of each individual, which was not provided in the dataset. To overcome this, we inferred each participant’s HLA genotype with a classifier that was based on previously published HLA-associated TCR β-chain sequences (DeWitt et al., 2018) and their abundance in each patient’s repertoire (see Methods for details). MIRA TCRs were not used to assign HLA-types to the 694 COVID-19 patients. We then used a beta-binomial counts regression model (Rytlewski et al., 2019) with each TCR feature to test for an association of feature abundance with presence of the restricting allele in the participant’s HLA genotype, controlling for participant age, sex, and days since COVID-19 diagnosis.

The models revealed that there were radius-defined meta-clonotypes with a strong positive and statistically significant association (FDR < 0.01) for 11 of the 17 HLA-restricted-MIRA sets that were evaluated (Figure 5A, Table S7); the significant HLA regression odds ratios ranged from 1.4 to 40 (median 4.9), indicating log-fold differences in meta-clonotype frequency between patients with and without the HLA genotype. Across all MIRA sets, an HLA-association (FDR < 0.01) was detected for 51.5% (943/1831) and 59.7% (830/1831) of the meta-clonotypes using the RADIUS or RADIUS+MOTIF definitions, respectively. In comparison, an HLA-association (FDR < 0.01) was detected for fewer than 3.7% (69/1831) of EXACT centroid features, largely because the specific TRBV gene and CDR3 sequences discovered in the MIRA experiments were infrequently observed in unenriched bulk samples (Figure 5B). When detectable, the abundance of centroid TCRs in bulk repertoires tended to be positively associated with expression of the restricting HLA allele, as hypothesized. However, in most cases, the associated false discovery rate-adjusted q-value of these associations were orders of magnitude larger (i.e., less statistically significant) than those obtained from using the engineered RADIUS or RADIUS+MOTIF feature with the same clonotype as a centroid (Figure 6B). The improved performance of meta-clonotypes as query features is particularly evident when testing for HLA-associated enrichment of TCRs recognizing immunodominant MIRA1 A*01, MIRA48 A*02, MIRA51 A*03, MIRA53 A*24, and MIRA55 A*01 (Figure 6). Moreover, the regression models with meta-clonotypes also revealed possible negative associations between TCR abundance and participant age and positive associations with sample collection more than two days post COVID-19 diagnosis (Figure 6A).

**Figure 5.**
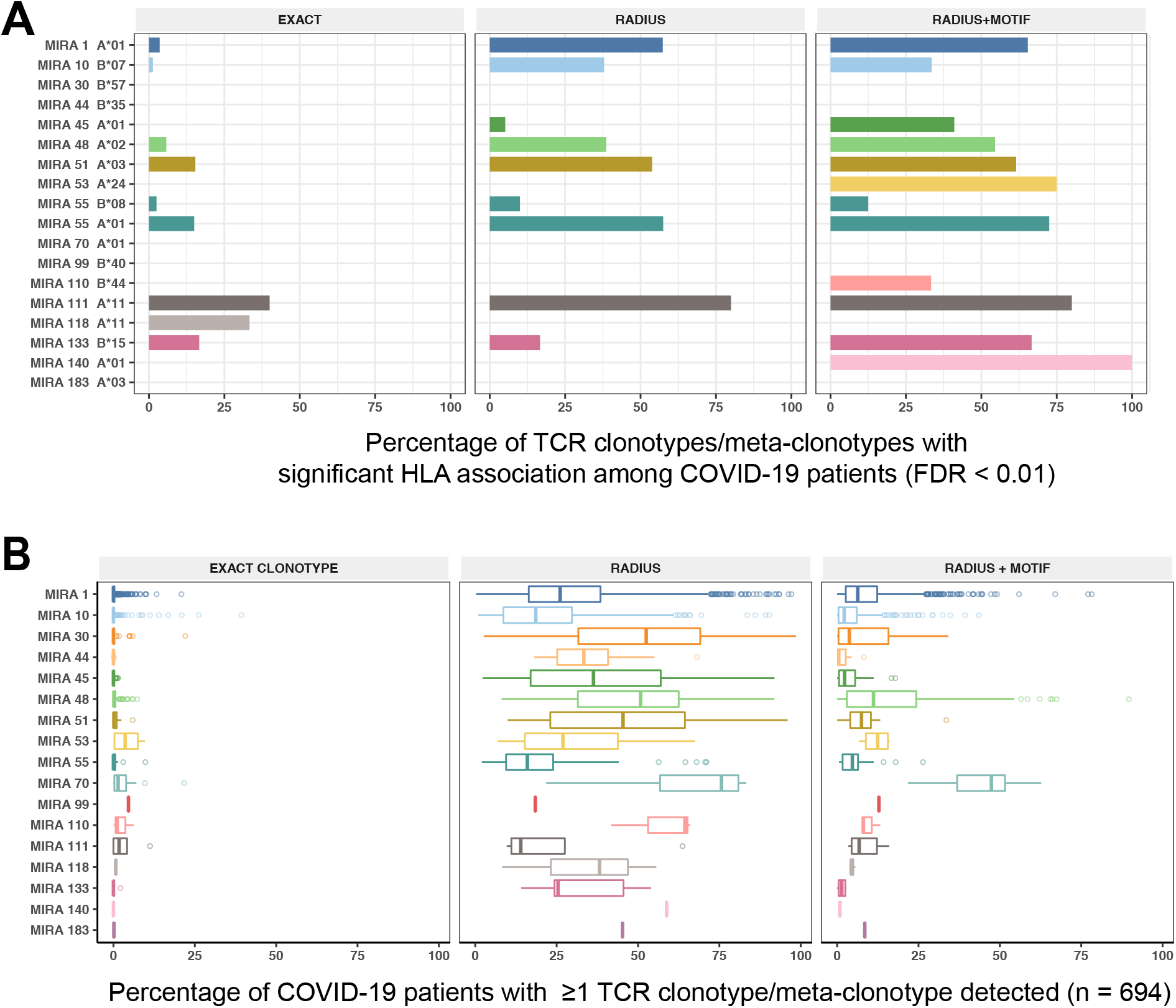
HLA restriction of TCR clonotypes and meta-clonotypes in bulk sequenced TCRβ repertoires of COVID-19 patients. (A) Percentage of TCR features with a statistically significant (FDR < 0.01) association with a restricting HLA allele. We tested for associations between patients’ inferred genotype and TCR feature abundance using beta-binomial regression controlling for age, sex, and days since COVID-19 diagnosis. (B) For each clonotype/meta-clonotype, the percent of bulk repertoires from COVID-19 patients (n=694) containing TCRs meeting the criteria defined by (1) EXACT (TCRs matching the centroid TRBV gene and amino acid sequence of the CDR3), (2) RADIUS (TCR centroid with inclusion criteria defined by an optimized TCRdist radius), or (3) RADIUS + MOTIF (inclusion criteria defined by TCR centroid, optimized radius, and the CDR3 motif constraint). See Figure 1 and Methods for details.

**Figure 6.**
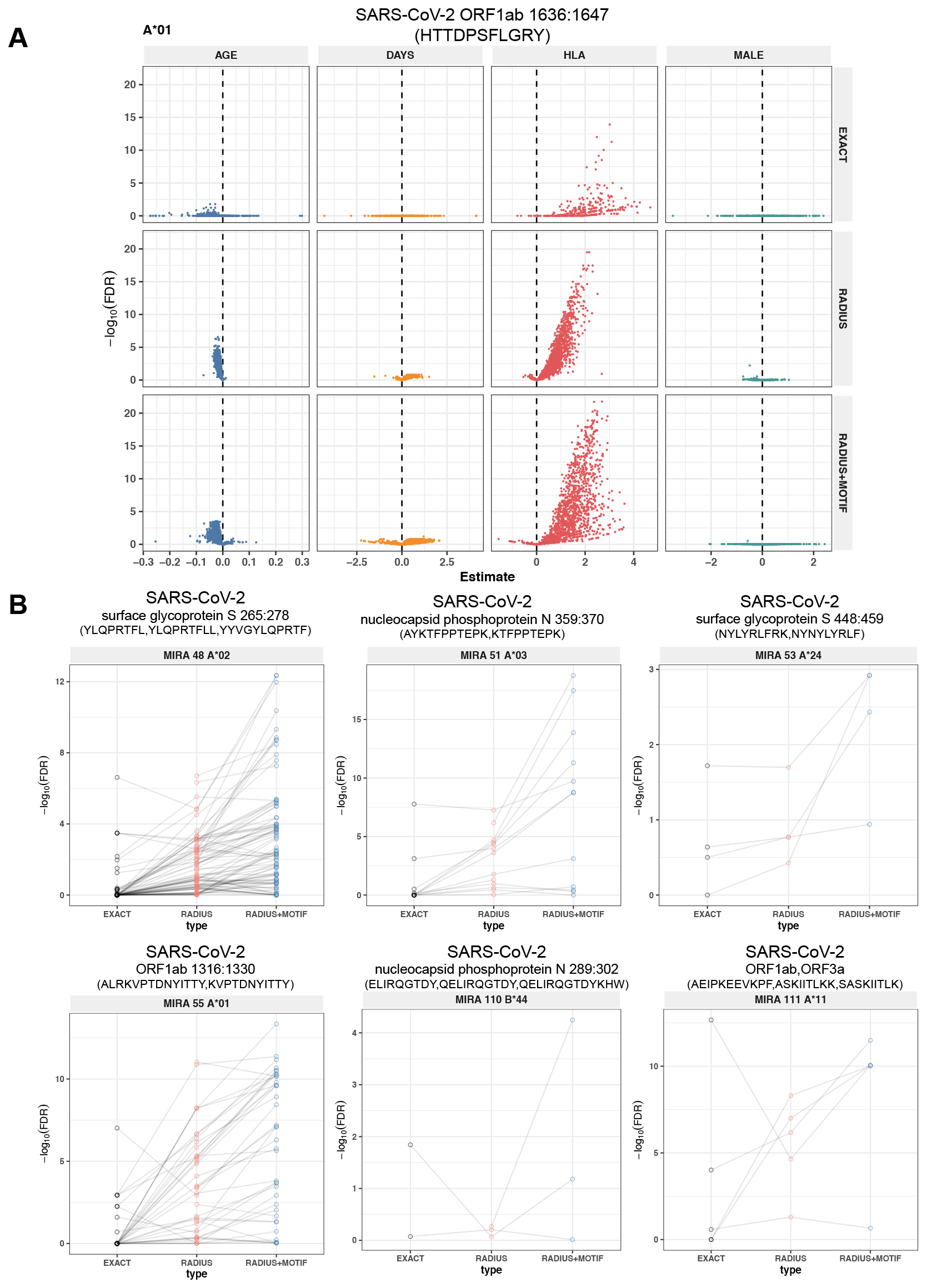
Associations of TCR features with participant age, days post diagnosis, HLA-genotype, and sex in TCR β-chain repertoires of COVID-19 patients (n=694). (A) Beta-binomial regression coefficient estimates (x-axis) and negative log_10_ false discovery rates (y-axis) for features developed from CD8+ TCRs activated by SARS-CoV-2 MIRA55 ORF1ab amino acids 1636:1647, HTTDPSFLGRY. The abundances of TCR meta-clonotypes are more robustly associated with predicted HLA type than exact clonotypes. (B) Signal strength of enrichment by participant HLA-type (2-digit) of TCR β-chain clonotypes (EXACT) and meta-clonotypes (RADIUS or RADIUS+MOTIF) predicted to recognize additional HLA-restricted SARS-CoV-2 peptides: (i) MIRA48 (ii) MIRA51 (iii) MIRA53 (iv) MIRA55 (v) MIRA110, and (vi) MIRA11 (See Table S6). Models were estimated with counts of productive TCRs matching clonotypes (EXACT) or meta-clonotypes (RADIUS or RADIUS+MOTIF) with the following definitions: (1) EXACT (inclusion of TCRs matching the centroid TRBV gene and amino acid sequence of the CDR3), (2) RADIUS (inclusion criteria defined by a TCR centroid and optimized TCRdist radius), (3) RADIUS + MOTIF (inclusion criteria defined by TCR centroid, optimized radius, and CDR3 motif constraint). See Methods for details.

### Comparison to k-mer based CDR3 features

Alternative methods exist for generating public TCR features from clustered clonotypes. One strategy is to identify clusters of TCRs that are each uniquely enriched with a short CDR3 k-mer, as implemented in GLIPH2 (Huang et al., 2020); this approach is well suited for identifying CDR3 k-mers associated with antigenic selection across bulk repertoires when knowledge of the specific antigens is unavailable (Chiou et al., 2021). To evaluate the similarities and differences of using this approach to generate public TCR features, compared with TCR distance-based meta-clonotypes, we applied tcrdist3 and GLIPH2 to 16 HLA-restricted MIRA sets (Figure 7; see Methods for details). Both methods identified public molecular patterns from MIRA TCRs (Figure S2) that were strongly HLA-associated in the large independent cohort of COVID-19 diagnosed patients (Figure 7). For this non-standard application of GLIPH2, we found that specificity groups based on global CDR3 k-mers (e.g., ‘SFRTD.YE’) tended to be more consistently HLA-associated than specificity groups based on local k-mers (e.g., ‘FRTD’). Compared to the GLIPH2 specificity groups based on global CDR3 kmers, meta-clonotypes tended to show similar or more evidence of HLA-association (i.e., smaller FDR values) (Figure 7). MIRA55:ORF1ab is an illustrative example; both the tcrdist3 meta-clonotypes GLIPH2 TCR groups were more strongly associated with the predicted A*01:01 HLA-restriction than exact clonotypes, supporting the general applicability of using antigen-enriched repertoires to create public features from otherwise private antigen-recognizing TCRs. Inspection of the meta-clonotypes and GLIPH2 groups showed that they were often overlapping, with meta-clonotypes subsuming multiple GLIPH2 groups. For example, A*01-associated meta-clonotype TRBV5-5*01+S.G[QE]G[AS]F[ST]DTQ (p-value 1E-12) fully overlaps several A*01-associated GLIPH2 patterns including S.GQGAFTDT (p-value 1E-12), QGAF (p-value 1E-11), and SLG.GAFTDT (p-value 1E-6). Similarly, the A*01-associated meta-clonotype TRBV28*01+S[RLMF][RK][ST]ND].YEQ (p-value 1E-13) covers 21 global GLIPH motifs including SFRTD.YE (p-value 1E-10), SLRTD.YE (p-value 1E-7), and SF.TDSYE (p-value 1E-4) (Table S9). These observations suggest that the motif-constraints of the meta-clonotypes were able to match a broader set of antigen-specific CDR3s compared to any one GLIPH2 specificity pattern, which may have helped boost detection sensitivity in the COVID-19 repertoires.

**Figure 7.**
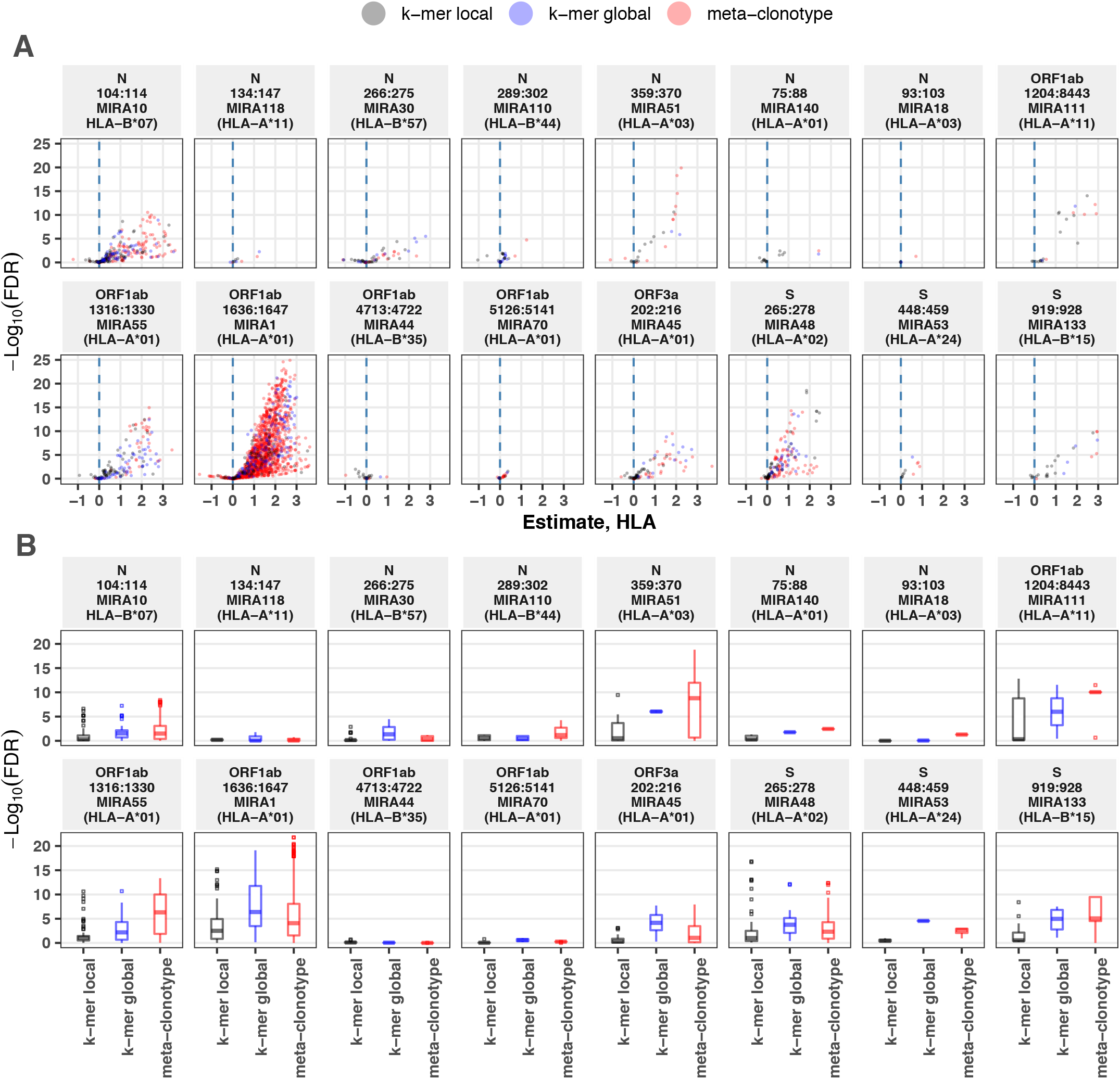
Associations between HLA-genotypes in COVID-19 patients and abundance of epitope specific CDR3 k-mers or meta-clonotypes. (A) Beta-binomial regression coefficient estimates (x-axis) for participant genotype matching a hypothesized restricting HLA allele and negative log_10_ false discovery rates (y-axis) for features developed from CD8+ TCRs activated by one of 16 HLA-restricted SARS-CoV-2 epitopes found in ORF1ab, ORF3a, nucleocapsid (N), and surface glycoprotein (S). Regression models included age, sex, and days post diagnosis as covariates (not shown). Positive HLA coefficient estimates correspond with greater abundance of the TCR feature in those patients expressing the restricting allele. (B) Distribution of false discovery rates by feature identification method (k-mer local, k-mer global, or meta-clonotype). Larger negative log10-tranformed FDR values (y-axis) indicate more statistically significant associations. Local k-mer (e.g., FRTD) and global k-mer (e.g., SFRTD.YE) were identified using GLIPH2 (Huang et al., 2020) and were used to quantify counts of conforming TCRs in each bulk sequenced COVID-19 repertoire (see Method for details).

## DISCUSSION

Given the extent of TCR diversity, only antigen-specific TCRs with high probability of generation (P_gen_) are likely to be detected reliably across individuals (Figure S3). While public, high-P_gen_ TCRs may sometimes be available for detecting a prior antigen-exposure, to more fully understand the population-level dynamics of complex polyclonal T-cell responses across a gradient of generation probabilities, it is critical to develop methods for finding public meta-clonotypes that capture otherwise private TCRs (Figure S3). We developed a novel framework, integrating antigen-enriched repertoires with efficiently sampled unenriched background repertoires, to engineer meta-clonotypes that balance the need for sufficiently public features with the need to maintain antigen specificity. The output of the analysis framework (Figure 1A) is a set of meta-clonotypes, each represented by a (i) centroid, (ii) radius, and (iii) optionally, a CDR3 motif-pattern, that can be used to rapidly search bulk repertoires for similar TCRs that likely share a cognate antigen. To demonstrate this analytical framework, we analyzed publicly available sets of antigen-enriched TCR β-chain sequences that putatively recognize SARS-CoV-2 peptides (Nolan et al., 2020). From these, we generated 4548 TCR radius-defined public meta-clonotypes that could be used to further investigate CD8+ T cell response to SARS-CoV-2 (Tables S7, S8).

To evaluate the potential relevance of radius-defined meta-clonotypes we focused on those with the strongest evidence of HLA restriction (Table S7, n=1831). We reasoned that we could compare the abundance of these meta-clonotypes in COVID-19 patients with and without the restricting HLA allele, where a significant positive association of abundance with expression of the restricting allele would provide confirmatory evidence both of the SARS-CoV-2 specificity of the meta-clonotype and its HLA restriction (Figure 1B). Overall, we found confirmation of HLA-restriction of meta-clonotype abundance for a majority of the MIRA sets we analyzed (11/17) and for 59% of meta-clonotypes tested using the RADIUS+MOTIF approach. There are several plausible explanations for the remaining meta-clonotypes that were not significantly HLA-restricted in this study. One possibility is that meta-clonotype definitions are not sufficiently specific for the target antigen; the radius is optimized for specificity, but not all amino acid substitutions accommodated within the radius are guaranteed to preserve antigen recognition, and while the motif constraint increases specificity, it’s likely that meta-clonotype definitions could be further refined with more antigen enriched TCR data and enhanced motif refinement methods. Also, sub-dominant SARS-CoV-2 epitopes may not be uniformly recognized, even among participants that share the required HLA genotype, which weakens the signal of HLA restriction detectable by regression analysis. A subset of GLIPH2 k-mer patterns were similar in their success at identifying groups of TCRs that confirmed the HLA restriction; it appeared that meta-clonotypes were generally more sensitive at the task, possibly afforded by non-conserved and multiple degenerate positions in the motif and lack of a constraint on the length of the CDR3, both of which enabled single meta-clonotypes to cover multiple GLIPH2 groups.

Recently, Snyder et al. (2020) analyzed 1521 bulk TCR β-chain repertoires from COVID-19 patients in the immuneRACE dataset and an additional 3500 (not yet publicly available) repertoires from healthy controls to identify public TCR β-chains that could be used to identify SARS-COV-2 infected individuals with high sensitivity and specificity. Their results show that with sufficient data it is possible to engineer highly performant TCR biomarkers of antigen exposure from exact clonotypes. We show that by leveraging antigen-enriched TCR repertoires it is possible to engineer radius-defined TCR meta-clonotypes from a relatively small group of COVID-19 diagnosed individuals (n=62; HLA-typed n=47) that are frequently detected in much larger independent cohorts. We propose that meta-clonotypes constitute a set of potential features that could be leveraged in developing TCR-based clinical biomarkers that go beyond detection of infection or exposure. For example, biomarkers predictive of infection, disease severity or vaccine protection may all require different TCR features. Statistical and machine learning tools could be employed to identify the meta-clonotypes and meta-clonotype combinations that carry the relevant clinical signal. Much like any biomarker study, to establish a TCR-based predictor of a particular outcome, the features must be measured among a sufficiently large cohort of individuals, with a sufficient mix of outcomes.

Though demonstrating HLA restriction of the SARS-CoV-2 meta-clonotypes establish their potential utility, it also highlighted how HLA diversity could be a major hurdle to biomarker development. The sensitivity of a TCR-based biomarker in a diverse population may depend on combining meta-clonotypes with diverse HLA restrictions since individuals with different HLA genotypes often target different epitopes using divergent TCRs. Our analysis shows that having HLA genotype information for TCR repertoire analysis can be critical to interpreting results. The simple HLA classifier we developed suggests that in the near future it may be possible to infer high-resolution HLA genotype from bulk TCR repertoires, but until then it is valuable to have sequenced-based HLA genotyping. In the absence of HLA genotype information, it may still be feasible to generate informative TCR meta-clonotypes. For example, a poly-antigenic TCR-enrichment strategy (i.e., peptide pools or whole-proteins) could help generate meta-clonotypes that broadly cover HLA diversity if the sample donors are racially, ethnically and geographically representative of the ultimate target population. For these reasons, donor unrestricted T cells and their receptors (e.g., MAITs, γδ T cells) may also be good targets for TCR biomarker development.

To enable TCR biomarker development and innovative extensions of distance-based immune repertoire analysis, we developed *tcrdist3*, which provides open-source (https://github.com/kmayerb/tcrdist3), documented (https://tcrdist3.readthedocs.io) computational building blocks for a wide array of TCR repertoire workflows in Python3. The software is highly flexible, allowing for: (i) customization of the distance metric with position and CDR-specific weights and amino acid substitution matrices, (ii) inclusion of CDRs beyond the CDR3, (iii) clustering based on single-chain or paired-chain data for α/β or γ/δ TCRs, and (iv) use of default as well as user-provided TCR repertoires as background for controlling meta-clonotype specificity (e.g., users may want to use HLA genotype-matched, or age-matched backgrounds). *tcrdist3* makes efficient use of available CPU and memory resources; as a reference, identification of meta-clonotypes from the MIRA55:ORF1ab dataset (n=479 TCRs) was completed in less than 5 minutes using 2 CPU and < 4 GB of memory including distance computation and radius optimization. Quantification of the identified meta-clonotypes (n=40) in 694 bulk TCR β-chain repertoires, ranging in size from 10,395 to 1,038,012 in-frame clones (~5 billion total pairwise comparisons) could be completed in less than 2 hours using 2 CPU and < 6 GB memory. The package also can generate multiple types of publication-ready figures (e.g., background-adjusted CDR3 sequence logos, V/J-gene usage chord diagrams, and annotated TCR dendrograms). The continued maturation of multiple adaptive immune receptor repertoire sequencing technologies will open possibilities for basic immunology and clinical applications, and *tcrdist3* provides a flexible tool that researchers can use to integrate the data sources needed to detect and quantify antigen-specific TCR features.

## METHODS

### TCR Data: immuneRACE datasets and MIRA assay

The study utilized two primary sources of TCR data (Nolan et al. 2020; Snyder et al. 2020). The first data source was a table of TCR β-chains amplified from CD8+ T cells activated after exposure to a pool of SARS-CoV-2 peptides, using a Multiplex Identification of Receptor Antigen (MIRA) (Klinger et al. 2015); data was accessed Jul 21, 2020 and labeled “ImmuneCODE-MIRA-Release002”. The samples used for the MIRA analysis included samples from 62 individuals diagnosed (3 acute, 1 non-acute, 58 convalescent) with COVID-19, of whom 47 (3 acute, 44 convalescent) were HLA-genotyped in the ImmuneCODE-MIRA-Release002 *subject-metadata.csv* file. We also used TCRs evaluated by MIRA from 26 COVID-19-negative control subjects that were part of ImmuneCODE-MIRA-Release002. We analyzed the 252 MIRA sets with at least 6 unique TCRs, referred to as MIRA1-MIRA252 in rank order by their size (Table S2); each “MIRA set” included antigen-associated TCRs across all assayed individuals. Adaptive Biotechnologies also made publicly available bulk unenriched TCR β-chain repertoires from COVID-19 patients participating in a collaborative immuneRACE network of international clinical trials. We selected repertoires from 694 individuals where meta-data was available indicating that the sample was collected from 0 to 30 days from the time of diagnosis. (COVID-19-DLS (Alabama, USA n = 374); COVID-19-HUniv12Oct (Madrid, Spain n = 117); COVID-19-NIH/NIAID (Pavia, Italy n=125) + COVID-19-ISB (Washington, USA n = 78). The sampling depth of these repertoires varied from 15,626-1,220,991 productive templates (median 208,709) and 10,395-1,038,012 productive rearrangements (median 113,716). We did not use bulk samples from the COVID-19-ADAPTIVE dataset as the average age was substantially lower than other immuneRACE populations and to avoid possible overlap with individuals that contributed samples to the MIRA experiments.

### HLA genotype inferences

No publicly available HLA genotyping was available for the 694 bulk unenriched immuneRACE T cell repertoires (Nolan et al. 2020). Before considering SARS-CoV-2 specific features, we inferred the HLA genotypes of these participants based on their TCR repertoires. Predictions were based on previously published HLA-associated TCR β-chain sequences (DeWitt et al., 2018) and their detection in each repertoire. Briefly, a weight-of-evidence classifier for each HLA loci was computed as follows: For each sample and for each common allele, the number of detected HLA-diagnostic TCR β-chains was divided by the total possible number of HLA-diagnostic TCR β-chains. The weights were normalized as a probability vector and the two highest HLA-allele probabilities (if the probability was larger than 0.2) were assigned to each repertoire; homozygosity was inferred if only one allele had probability > 0.2. The sensitivity and specificity of this simple classifier for each allele prediction were assessed using 550 HLA-typed bulk repertoires (Emerson et al., 2017). Sensitivities for common HLA-A alleles A*01:01, A*02:01, A*03:01, A*24:02, and A*11:01 were 0.96, 0.91, 0.90. 0.84, 0.94, respectively. Specificity for major HLA-A alleles was between 0.97-1.0. Inference of the HLA genotype of most participants was deemed sufficient in the absence of direct HLA genotyping.

### Peptide-HLA binding prediction

HLA binding affinities of peptides used in the MIRA stimulation assay were computationally predicted using NetMHCpan4.0 (Jurtz et al., 2017). Specifically, the affinities of all 8, 9, 10 and 11mer peptides derived from the stimulation peptides were computed with each of the class I HLA alleles expressed by participants in the MIRA cohort (n=47). From this data we derived 2-digit HLA binding predictions (e.g., A*02) for each MIRA set by pooling the predictions for all the 4-digit HLA variants (e.g. A*02:01, A*02:02) across all the derivative peptides and selecting the lowest IC50 (strongest affinity). Predictions with IC50 < 50 nM were considered strong binders and IC50 < 500 nM were considered weak binders (Table S3, Table S4).

### TCR distances

Weighted multi-CDR distances between TCRs were computed using *tcrdist3*, an open-source Python3 package for TCR repertoire analysis and visualization, using the procedure first described in (Dash et al., 2017). The package has been expanded to accommodate γδ TCRs; it has also been recoded to increase CPU efficiency using *numba*, a high-performance just-in-time compiler. A numba-coded edit/Levenshtein distance is also included for comparison, with the flexibility to accommodate novel TCR metrics as they are developed.

Briefly, the distance metric in this study is based on comparing TCR β-chain sequences. The *tcrdist3* default settings compare TCRs at the CDR1, CDR2, and CDR2.5 and CDR3 positions. By default, IMGT aligned CDR1, CDR2, and CDR2.5 amino acids are inferred from TRVB gene names, using the *01 allele sequences when allele level information is not available. The CDR3 junction sequences are trimmed 3 amino acids on the N-terminal side and 2 amino acids on the C-terminus, positions that are highly conserved and less crucial for mediation of antigen recognition. For two CDR3s with different lengths, a set of consecutive gaps are inserted at a position in the shorter sequence that minimizes the summed substitution penalties based on a BLOSUM62 substitution matrix. Distances are then the weighted sum of substitution penalties across all CDRs, with the CDR3 penalty weighted three times that of the other CDRs.

### Optimized TCR-specific radius

To find biochemically similar TCRs while maintaining a high level of specificity, we used the packages *tcrdist3* and *tcrsampler* to generate an appropriate set of unenriched antigen-naïve background TCRs. A background repertoire was created for each MIRA set, with each consisting of two parts. First, we combined a set of 100,000 synthetic TCRs generated using the software OLGA (Marcou et al., 2018; Sethna et al., 2019), whose TRBV- and TRBJ-gene frequencies match those in the antigen-enriched repertoire. Second we used 100,000 umbilical cord blood TCRs sampled evenly from 8 subjects (Britanova et al., 2016). This mix balances dense sampling of background sequences near the biochemical neighborhoods of interest with broad sampling of common TCR representative of antigen-naive repertoire. We then adjust for the biased sampling by using the TRBV- and TRBJ-gene frequencies observed in the cord-blood data. The adjustment is a weighting based on the inverse of each TCR’s sampling probability. Because we oversampled regions of the “TCR space” near the candidate centroids we were able to estimate the density of the meta-clonotype neighborhoods well below 1 in 200,000. This is important because ideal meta-clonotypes would be highly specific even in repertoires larger than 200,000 sequences. With each candidate centroid, a meta-clonotype was engineered by selecting the maximum distance radius that still controlled the number of neighboring TCRs in the weighted unenriched background to 1 in 10^6^. Candidate centroids that used a TRBV gene that was not in the cord-blood repertoires were excluded from further analysis, since an estimate of gene frequency is required to apply the inverse weighting described above.

### TCR meta-clonotype MOTIF constraint

Radius-optimized meta-clonotypes from antigen-enriched TCRs-provided an opportunity to discover key conserved residues most likely mediating antigen specificity. We developed a “motif” constraint as an optional part of each meta-clonotype definition that limited allowable amino-acid substitutions in highly conserved positions of the CDR3 to those observed in the antigen-enriched TCRs. The motif constraint for each radius-defined meta-clonotype was defined by aligning each of the conformant CDR3 amino-acid sequences to the centroid CDR3. Alignment positions with five or fewer distinct amino acids were considered conserved and added to the motif as a set of possible residues. Thus, the motif constraint is permissive of only specific substitutions in select positions relative to the centroid, however these substitutions are still penalized by the radius constraint. The motif constraint was encoded as a regular expression, with the “.” character indicating non-conserved positions and bracketed residues indicating a degenerate position with a set of allowable residues (e.g., “SL[RK][ND]YEQ”). Position with gaps, where some sequences are missing a residue, are accommodated by making that position optional (e.g., “SL[RK]?[ND]YEQ”). Since the motif constraints form regular expressions, they can be used to rapidly scan large repertoires for conformant CRs and easily be combined with a radius constraint. When applied to bulk repertoires, the motif constraint eliminates CDR3s that did not match key conserved residues.

### TCR abundance regression modeling

Similar to bulk RNA sequencing data, TCR frequencies are count data drawn from samples of heterogeneous size. Thus we initially attempted to fit a negative binomial model to the data (e.g., DESEQ2 (Love et al., 2013)). We found that the negative binomial model did not adequately fit TCR counts, which – compared to transcriptomic data – were characterized by (i) more technical zeros due to inevitable under sampling and (ii) even greater biological over-dispersion, which could be due to clonal expansions and HLA genotype diversity. Instead we found that the beta-binomial distribution, which was recently used for TCR abundance modeling (Rytlewski et al., 2019), provided the flexibility needed to adequately fit the TCR data. We used an R package, *corncob*, which provides maximum likelihood methods for inference and hypothesis testing with beta-binomial regression models (Martin et al., 2020). Due to the sparsity of some meta-clonotypes, seven percent of coefficient estimates in regression models had p-values larger than 0.99 (i.e., non-significant) and unreliable high magnitude coefficient estimates. These values are not shown in the horizontal range of the volcano plots. From the p-values for each regression coefficient we computed false-discovery rate (FDR) adjusted q-values and accepted q-values < 0.01 (1%) as statistically significant; adjustment was performed across meta-clonotypes within each MIRA set and within each variable class (e.g., HLA, age, sex, or days since diagnosis). The HLA regression coefficients from the beta-binomial models indicate log-fold differences in meta-clonotype abundance between patients with and without the HLA genotype.

### Comparison with k-mer based CDR3 features

*GLIPH2* (Huang et al., 2020) software *irtools.osx* was applied to 16 antigen-enriched sub-repertoire of TCRs with epitopes with strong prior evidence of restriction to an HLA-A or HLA-B allele to demonstrate how a k-mer based tool might also be used to cluster biochemically similar antigen-specific TCRs to discover potential TCR biomarker features. GLIPH2 generates “global” TCR specificity groups of CDR3s of identical length with a single optional non-conserved position based on enrichment frequency of ‘local’ continuous 2-mer, 3-mers, and 4-mers. We used the GLIPH2-provided ‘ref_CD8_v2.0.txt’ background file as a background to identify enriched features. Across epitope-specific MIRA sets, we tested HLA-associations of 812 GLIPH2 pattern ranging from 3 to 11 amino acids in length. The MIRA55:ORF1ab set was chosen for detailed analysis because, among the MIRA sets, it is comprised of CD8+ TCR β-chains activated by a peptide with the strongest evidence of HLA-restriction, primarily HLA-A*01. The MIRA55 set of TCRs, GLIPH2 returned 121 testable public clusters (based on 67 local k-mers, 54 global k-mers) associated with CDR3 patterns enriched relative the program’s default CD8+ TCR background (GLIPH2 default Fisher’s exact test, p-value < 0.001). The GLIPH2 patterns and their associated “specifity group” TRBV gene usages and sequence length were then used to search for conforming TCRs in the 694 bulk unenriched COVID-19 repertoires, allowing comparison to exact and meta-clonotype features. GLIPH2 represents degenerate positions using the “%” character, which we represent throughout this study by the “.” character.

### tcrdist3: Software for TCR repertoire analysis

*tcrdist3* is an open-source Python3 package for TCR repertoire analysis and visualization. The core of the package is the TCRdist, a distance metric for relating two TCRs, which has been expanded beyond what was previously published (Dash et al., 2017) to include γδ-TCRs. It has also been recoded to increase CPU efficiency using *numba*, a high-performance just-in-time compiler. A numba-coded edit/Levenshtein distance is also included for comparison, with the flexibility to accommodate novel TCR metrics as they are developed. The package can accommodate data in standardized format including AIRR, vdjdb exports, MIXCR output, 10x Cell Ranger output or Adaptive Biotechnologies immunoSeq output. The package is well documented including examples and tutorials, with source code available on github.com under an MIT license (http://github.com/kmayerbl/tcrdist3). *tcrdist3* imports modules from several other open-source, pip installable packages by the same authors that support the functionality of *tcrdist3*, while also providing more general utility. Briefly, the novel features of these packages and their relevance for TCR repertoire analysis is described here:

*pwseqdist* enables fast and flexible computation of pairwise sequence-based distances using either *numba*-enabled tcrdist and edit distances or any user-coded Python3 metric to relate TCRs; it can also accommodate computation of “rectangular” pairwise matrices: distances between a relatively small set of TCRs with all TCRs in a much larger set (e.g., bulk repertoire). On a modern laptop distances can be computed at a rate of ~70M per minute, per CPU.

*tcrsampler* is a tool for sub-sampling large bulk datasets to estimate the frequency of TCRs and TCR neighborhoods in non-antigen-enriched background repertoires. The module comes with large, bulk sequenced, default databases for human TCR α, β, γ and δ and mouse TCR β (Britanova et al., 2016; Ravens et al., 2018; Wirasinha et al., 2018). Datasets were selected because they represented the largest pre-antigen exposure TCR repertoires available; users can optionally supply their own background repertoires when applicable. An important feature of *tcrsampler* is the ability to specify sampling strata; for example, sampling is stratified on individual by default so that results are not biased by on individual with deeper sequencing. Sampling can also be stratified on V and/or J-gene usage to over-sample TCRs that are somewhat similar to the TCR neighborhood of interest. This greatly improves sampling efficiency, since comparing a TCR neighborhood to a background set of completely unrelated TCRs is computationally inefficient; however, we note that it is important to adjust for biased sampling approaches via inverse probability weighting to estimate the frequency of oversampled TCRs in a bulk-sequenced repertoire.

*palmotif* is a collection of functions for computing symbol heights for sequence logo plots and rendering them as SVG graphics for integration with interactive HTML visualizations or print publication. Much of the computation is based on existing methods that use either KL-divergence/entropy or odds-ratio based approaches to calculate symbol heights. We contribute a novel method for creating a logo from CDR3s with varying lengths. The target sequences are first globally aligned (parasail C++ implementation of Needleman-Wunsch) to a pre-selected centroid sequence (Daily, 2016). For logos expressing relative symbol frequency, background sequences are also aligned to the centroid. Logo computation then proceeds as usual, estimating the relative entropy between target and background sequences at each position in the alignment and the contribution of each symbol. Gaps introduced in the centroid sequence are ignored, while gap symbols in the aligned sequences are treated as an additional symbol.

## SUPPLEMENTAL TABLES

Table S1 Comparison of selected software tools for clustering TCRs

Table S2 MIRA enriched repertoires MIRA0 - MIRA252

Table S3 HLA class I alleles capable of presenting the SARS-CoV-2 associated peptides in MIRA screen

Table S4 NetMHCpan4.0 peptide MHC class I binding affinity prediction

Table S5 Statistical associations between common HLA genotypes of COVID-19 exposed MIRA participants and SARS-CoV-2 peptide-enriched TCR repertoires

Table S6 SARS-CoV-2 CD8+ meta clonotypes summarized by MIRA enriched repertoire

Table S7 SARS-CoV-2 CD8+ meta clonotypes with strong evidence of HLA restriction (n = 1831)

Table S8 SARS-CoV-2 CD8+ meta clonotypes with less evidence of HLA restriction (n = 2717)

Table S9 HLA associations of GLIPH2 k-mers and tcrdist3 meta-clonotypes

## SUPPLEMENTAL FIGURES

**Figure S1:**
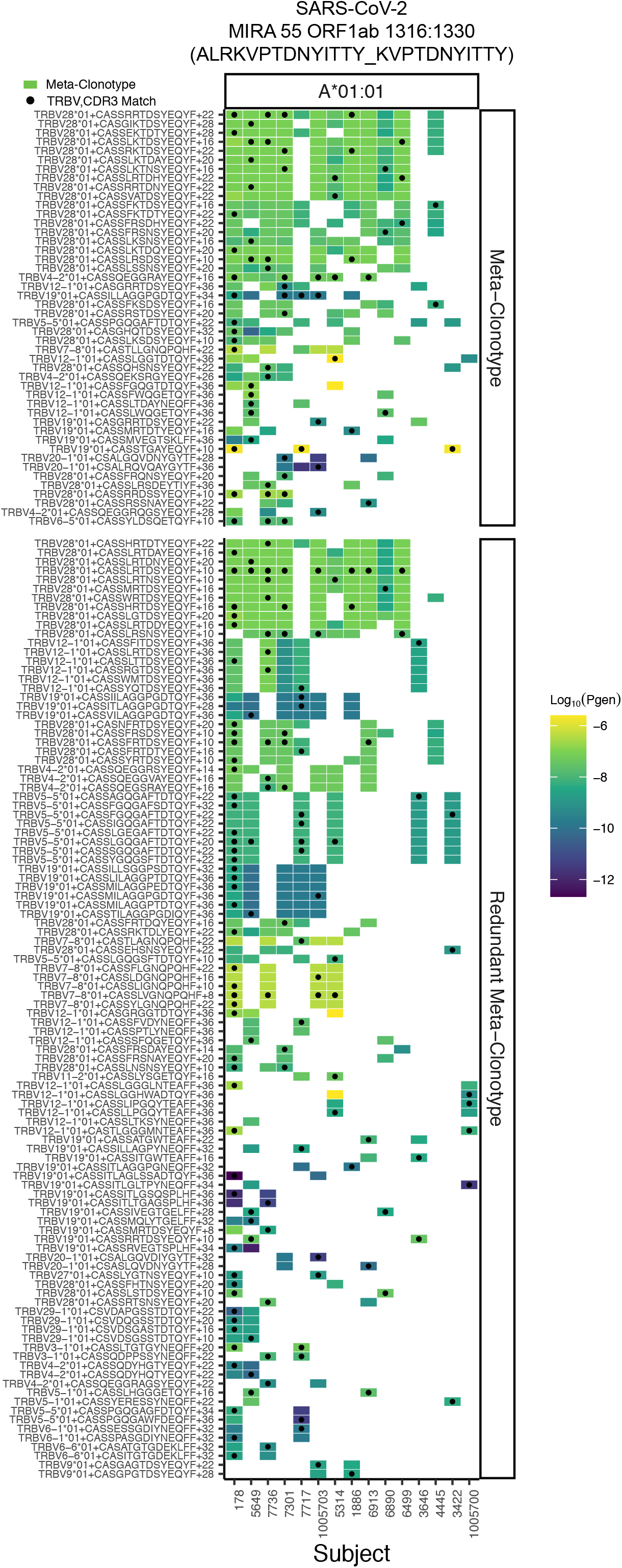
Publicity analysis in MIRA participants of CD8+ TCR β-chain features activated by SARS-CoV-2 peptide ORF1ab (MIRA55) predicted to bind HLA-A*01. The grid shows all features that were present in 2 or more MIRA participants. TCR feature publicity across individuals was assessed using two methods: (i) tcrdist3 *meta-clonotypes* (rectangles) – inclusion criteria defined by a centroid TCR and all TCRs within an optimized TCRdist radius selected to span < 10^−6^ TCRs in a bulk unenriched background repertoire, and (ii) exact public clonotypes (circles) are defined by matching TRBV gene usage and identical CDR3 amino acid sequence. Per subject, the color-scale shows the meta-clonotype conformant clone with the highest probability of generation (P_gen_). All TCRs captured by a “redundant” meta-clonotypes were completely captured by a higher ranked meta-clonotype. Redundant meta-clonotypes were not subsequently evaluated.

**Figure S2:**
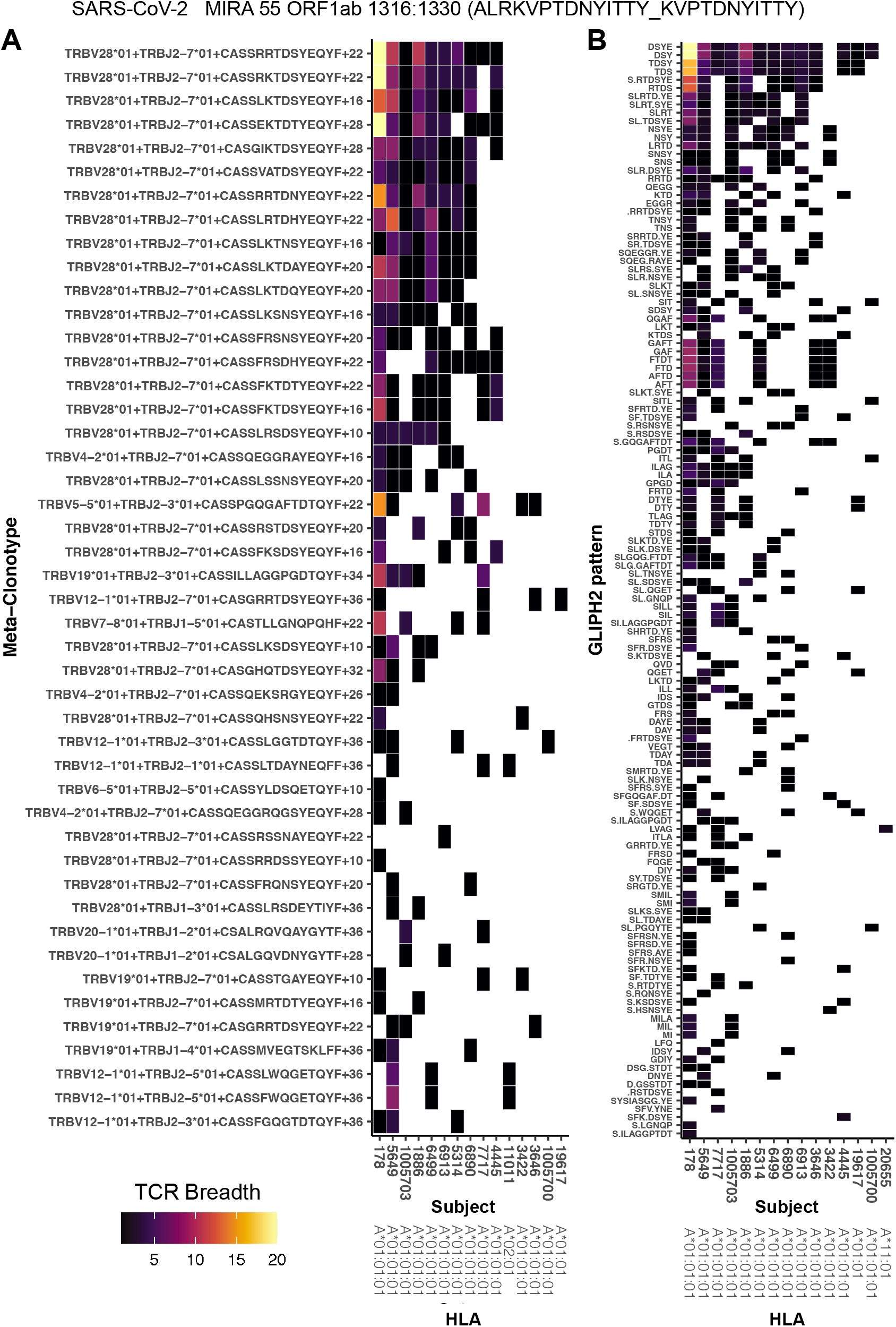
Publicity and breadth analysis of CD8+ TCR β-chain features activated by SARS-CoV-2 peptide ORF1ab (MIRA55) using *tcrdist3* and GLIPH2. TCR feature publicity was determined using two methods for clustering similar TCR sequences: (A) *tcrdist3*-identified meta-clonotypes and (B) GLIPH2 specificity-groups, sets of TCRs with a shared CDR3 k-mer pattern uncommon in the program’s default background CD8+ receptor data. Grid fill color shows the breadth – or number of conformant clones – withing each patient’s repertoire.

**Figure S3:**
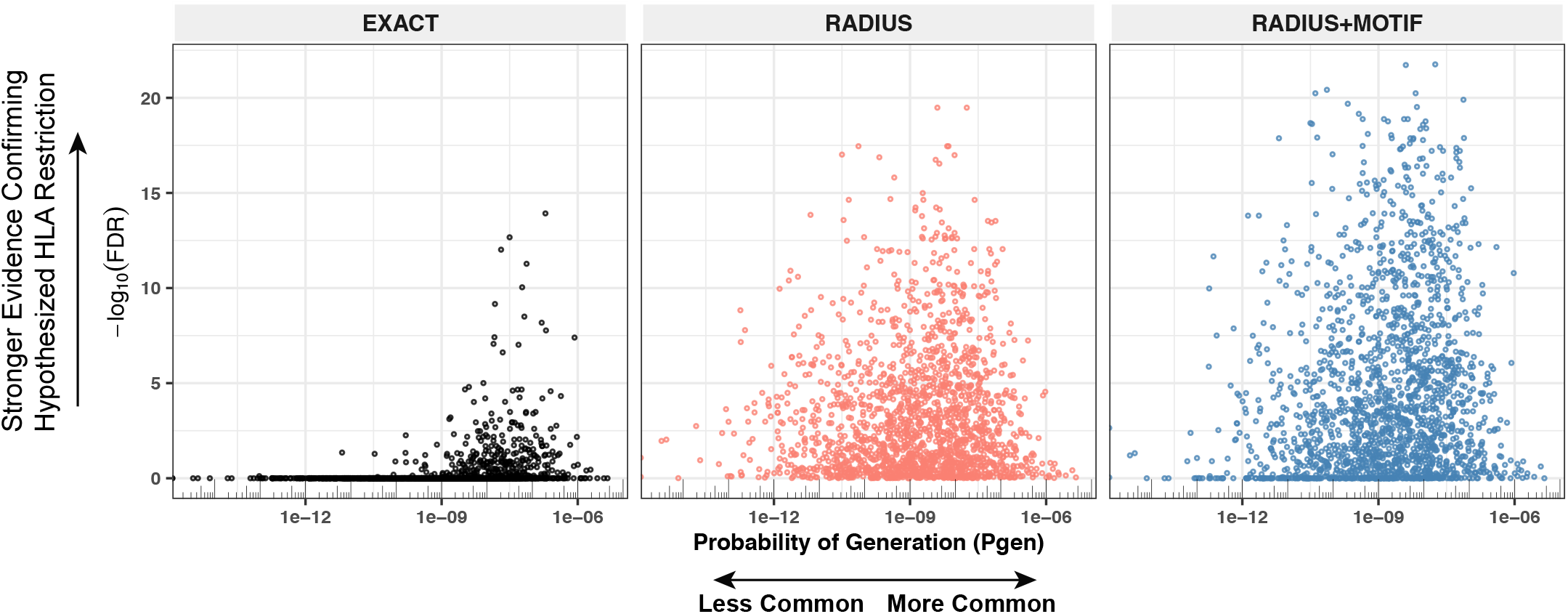
Detectable HLA-association and CDR3 probability of generation. We evaluated meta-clonotypes from 17 MIRA sets in a cohort of 694 COVID-19 patients for their association with predicted HLA-restricting alleles. Statistical evidence of the HLA association for each meta-clonotype (RADIUS or RADIUS+MOTIF) and the centroid alone (EXACT) is indicated by the associated false discovery rate (FDR; y-axis) in beta-binomial regressions (see Methods for model details). The probability of generation (Pg en) of each centroid’s CDR3-β was estimated using the software OLGA (x-axis). Using exact matching, only associations with high probability of generation (P_gen_) antigen-specific TCRs are likely to be detected reliably. However, using meta-clonotypes, *tcrdist3* revealed strong evidence of HLA-restriction for TCRs with both high and low probability of generation.

## DATA AVAILABILITY

ImmuneRACE data is publicly available: https://immunerace.adaptivebiotech.com/data/. All other TCR data is publicly available from VDJdb (https://vdjdb.cdr3.net/) or the cited research.

## SOFTWARE AVAILABILITY

The *tcrdist3* code base used in this analysis is freely available at https://github.com/kmayerb/tcrdist3/ with documented examples at http://tcrdist3.readthedocs.io. *tcrdist3* relies on the Python package *pwseqdist* - freely available at https://github.com/agartland/pwseqdist - for numba-optimized just-in-time compiled versions of the TCRdist measure.

## CONTRIBUTIONS

Conceptualization: KM, SS, LC, JCC, AS, JG, TH, PT, PB, AF; Methodology; Software: KM, AF; Validation; Formal analysis; Investigation: KM, AF; Data Curation; Writing – original draft preparation: KM, AF; Writing – review & editing: KM, SS, LC, JCC, AS, JG, TH, PT, PB, AF; Supervision: TH, PT, PB, AF; Funding acquisition: TH, PT, PB, AF, JCC

## ACKNOWLEDGEMENTS

This work was funded by NIH NIAID R01 AI136514-03 (PI Thomas) and ALSAC at St. Jude. The authors thank M. Pogorelyy and A. Minervina for extensive feedback on the manuscript. Scientific Computing Infrastructure at Fred Hutchinson Cancer Research Center was funded by ORIP grant S10OD028685.

## Notes

### Competing Interest Statement

PT is on the Scientific Advisory Boards of Immunoscape and Cytoagents, consulted for Elevate Bio and PACT Pharma, and has received travel costs and speaking fees from 10X Genomics and Illumina. PT also has filed patents on methods for sequencing and cloning TCRs. PB, PGT, and JCC served as unpaid consultants for 10X Genomics on the initial analysis of the 10x_200k dataset. TH has equity in Poold Diagnostics.

### Summary of Updates

Main text updated, Figure 7 added, and Supplemental Tables updated.

